# Distinct microcolony morphologies promote flow-dependent bacterial colonization

**DOI:** 10.1101/2023.11.22.568348

**Authors:** Kelsey M. Hallinen, Steven P. Bodine, Howard A. Stone, Tom W. Muir, Ned S. Wingreen, Zemer Gitai

## Abstract

Fluid flows can impact bacterial behaviors in unexpected ways (*1*–*3*). The high shear rate in heart valves should reduce colonization, but in endocarditis, valves are often counter-intuitively colonized by *Staphylococcus aureus* and *Enterococcus faecalis* (*4, 5*). Here we discover bacteria-specific mechanisms for preferential surface colonization in higher shear rate environments. This behavior enables bacteria that are outcompeted in low flow to dominate in high flow. Flow-dependent colonization by *S. aureus* and *E. faecalis* are mediated by distinct mechanisms that depend on each species’ microcolony morphologies: transport of a dispersal signaling molecule for clustered *S. aureus* and mechanical forces for linear chains of *E. faecalis*. These results suggest that microcolony morphologies have previously unappreciated costs and benefits in different environments, like those introduced by flow.

**One-Sentence Summary:** Bacterial surface colonization in high fluid flow depends upon the species’ clustered or chained microcolony morphologies.

## Introduction

Bacterial populations experience a range of complex environments in their natural settings (*2*). For example, within a host, bacteria must contend with fluid flows such as those found in the cardiovascular system (*6*). Fluid flow can significantly impact how bacteria interact with one another or with the surfaces that they colonize (*7*–*13*). Recent studies have begun to examine the effects of flow on single-cell bacterial behaviors like growth (*14, 15*) and adhesion(*16*–*21*). For example, catch bonds can enable individual bacterial cells to adhere to epithelial surfaces more strongly in flow(*22*). However, these single-cell mechanisms are often not widely conserved. Furthermore, in natural settings bacteria are typically in dynamic multicellular communities and the effects of flow on cell collectives are underexplored.

One example of a counterintuitive bacterial behavior in the presence of flow comes from clinical reports of infective endocarditis. Infective endocarditis is an infection in the heart, where bacteria are most commonly reported to colonize heart valves (*4, 5, 20, 23, 24*). These infections are difficult to treat and even with the use of antibiotics surgery is often needed to replace infected valves (*25*). From the perspective of fluid mechanics, the valves have the narrowest cross-sectional areas in the heart, causing them to have the fastest flow speed (*6*). These observations lead to an intriguing paradox: higher flow speeds would naively be thought to reduce surface colonization, so how might bacteria preferentially colonize the regions of the heart with the highest flow conditions?

Here we sought to directly examine the effects of flow on the surface colonization of bacteria that cause endocarditis. To this end, we studied two different bacterial species commonly implicated in endocarditis infections, Methicillin-resistant *Staphylococcus aureus* (MRSA) and *Enterococcus faecalis*, and imaged their surface colonization within microfluidic devices. Our results demonstrate that these bacteria preferentially colonize surfaces in high shear conditions, even when the surface is abiotic, suggesting that this counterintuitive behavior is driven by the bacteria themselves. Elucidating how the two species preferentially adhere in high shear environments reveals two different mechanisms that rely on their distinct microcolony morphologies and are driven by either transport of signaling molecules or mechanical responses to flow.

## Results

### Bacterial populations exhibit counterintuitive adhesion behaviors in microfluidic studies

To assess the impact of flow on bacteria in the absence of host factors, we examined the colonization of glass coverslips for both *S. aureus* (USA300 MRSA) and *E. faecalis* (OG1RF) in untreated microfluidic channels (Fig. 1A). For all experiments described below, cells were seeded into the channel and allowed to adhere to the glass coverslip before we flowed in sterile media, such that no new cells were introduced to the system once flow began. In any given flow system, there is a flow rate (volume/time) and a typical speed (flow rate divided by the cross-sectional area of the flow). The fluid speed is zero at the boundaries, so cells at the surface typically experience a flow (or velocity) profile that increases with distance from the wall. Hence, the cells attached to a wall experience a shear rate (units 1/second). For a given configuration, higher flow rates correspond to higher shear rates. The shear rate in heart valves is estimated to be roughly 100/s (estimated from (*26, 27*) as detailed in the supplement). For each species, we thus initially examined two shear rates: a low shear rate of 40/s (low flow) and a high shear rate of 400/s (high flow). Despite the naive expectation that higher shear rate should hinder colonization, by the end of our experiments we observed that more bacteria colonized the surface for both *S. aureus* and *E. faecalis* (Fig. 1). Performing similar experiments on other bacterial species like *Escherichia coli*, and *Streptococcus pneumoniae* revealed that these species do not increase colonization in high flow, suggesting that this behavior is specific to certain species (Movie S1-S4, Fig. S1).

**Fig. 1.**
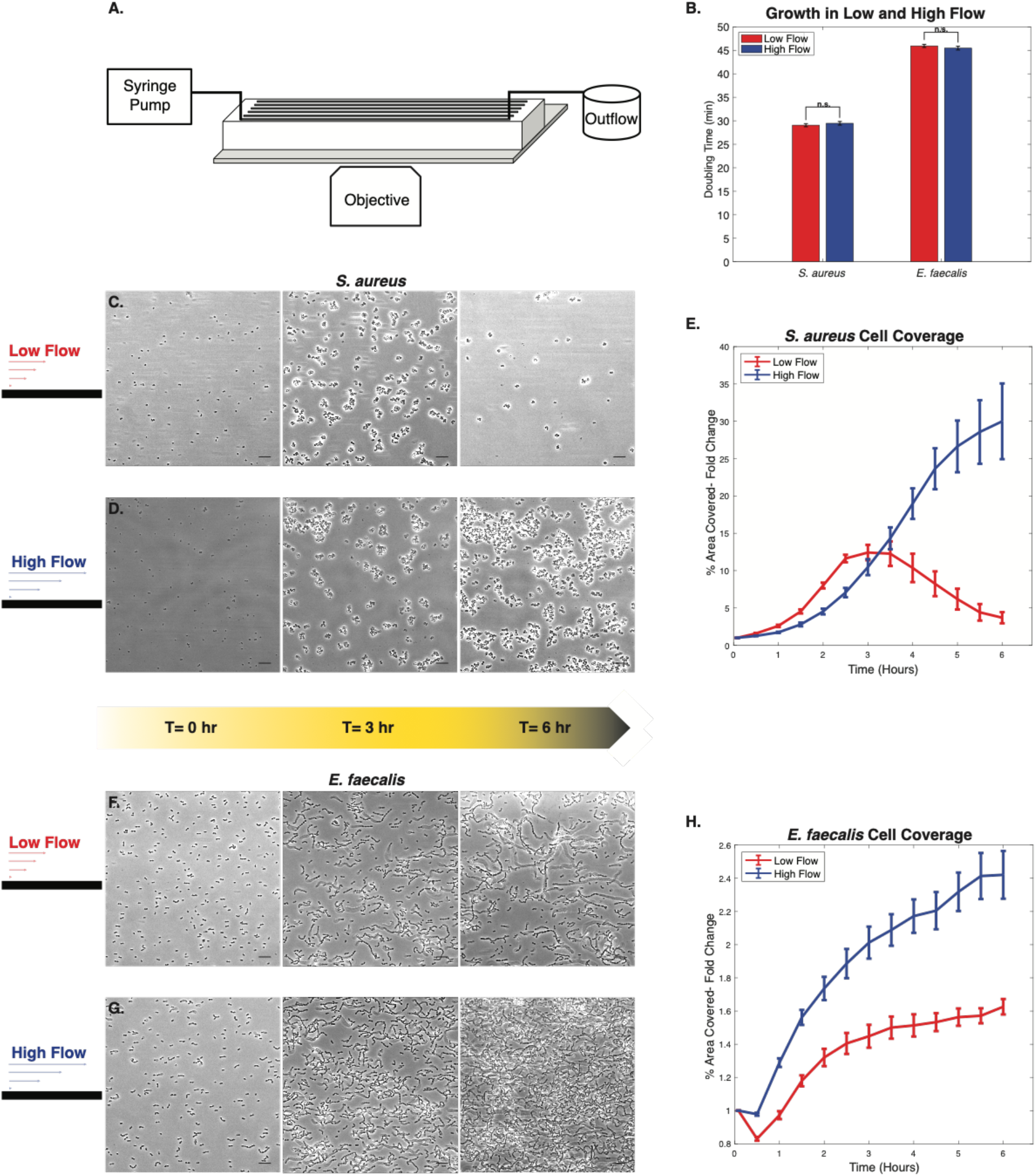
Paradoxical final cell population in low versus high flow. **(A)** Schematic of microfluidic chamber used for all flow experiments. A PDMS microfluidic channel is bonded to a glass coverslip. Following cell loading, a syringe pump is turned on, flowing fresh media through the channel. Cells attached to the coverslip are imaged. (B) Analysis of doubling times for each *S. aureus* MRSA and *E. faecalis* in low (red) and high (blue) flow conditions. (C) Representative images of *S. aureus* at 0, 3, and 6 hours of low flow (shear rate 40/s). (D) Similar representative images of *S. aureus* at 0, 3, and 6 hours of high flow (shear rate 400/s). (E) Fold change of percent area covered of both low (red) and high (blue) flow experiments for *S. aureus*. (F) Representative images of *E. faecalis* at 0, 3, and 6 hours of low flow (shear rate 40/s). (G) Similar representative images of *E. faecalis* at 0, 3, and 6 hours of high flow (shear rate 400/s). (H) Fold change of percent area covered of both low (red) and high (blue) flow experiments for *E. faecalis*. Image scale bars are 10 μm. Error bars are SEM.

Closer examination of our timelapse movies of *S. aureus* and *E. faecalis* indicated that the attachment of new cells to the surface was relatively rare in both high flow and low flow for both species (Movie S5-8). Furthermore, analysis of single cell doubling time showed no significant difference between the low and high flow conditions for either species, indicating that the differences in colonization cannot be explained by differences in growth rate (Fig. 1B). Since the differences between high and low flow could not be explained by differences in attachment or growth, we focused our subsequent studies on understanding mechanisms driving differences in detachment.

To better understand flow-dependent *S. aureus* colonization, we examined colonization dynamics and found that for the first several hours surface colonization increased at a similar rate in both low and high flow. During these early timepoints the bacteria formed small microcolonies with a clustered morphology. After roughly 3 hours, however, a striking difference between the low and high flow conditions began to emerge. At these later timepoints, *S. aureus* continued to colonize more and more of the surface in the high flow conditions. But in the low flow condition, *S. aureus* began to disperse from the surface, leading to a decrease in coverage and smaller microcolony clusters. Image analysis of the fraction of the surface area covered by bacteria confirmed that in high flow colonization proceeded to monotonically increase, but that in low flow, colonization increased for the first 3 hours and then exhibited increasing dispersal for the rest of the experiment (Fig. 1C-E).

The dynamics of *E. faecalis* colonization revealed a different pattern from that of *S. aureus*. Throughout the experiments we found that *E. faecalis* grew as microcolonies with a linear chained morphology (Fig. 1F-G). In both low and high flow, *E. faecalis* surface colonization increased mostly monotonically, but the rate of the increased surface colonization was consistently higher in high flow than in low flow conditions (Fig. 1F-H). Together, these experiments suggest that the surface colonization of both *E. faecalis* and *S. aureus* is counterintuitively greater in high flow than in low flow. While changes in detachment rate appear to underlie both behaviors, the two species respond to flow differently, as they exhibit distinct colonization dynamics.

### Flow-dependent competition dynamics between *S. aureus* and *P. aeruginosa*

Could preferential surface colonization provide a competitive advantage in flow? *S. aureus* and *Pseudomonas aeruginosa* are two bacterial pathogens that are often found together in polymicrobial infections. Previous studies have established that in the absence of flow *P. aeruginosa* generally outcompetes *S. aureus* (*28*). However, the interactions between these species have not been previously examined in higher-flow environments. We thus mixed *S. aureus* and *P. aeruginosa* and examined their surface colonization dynamics in our system. We first studied their dynamics in a minimal flow condition (shear rate of 2/s), to keep the channel clear and provide fresh media. As expected, in these co-cultures *P. aeruginosa* outcompeted *S. aureus* and quickly came to dominate the channel (Fig. 2A,B, Movie S9). In contrast, observing the co-cultures in high flow conditions (shear rate of 400/s) showed the opposite effect. In these conditions, *S. aureus* robustly colonized the surface and outcompeted *P. aeruginosa* to ultimately dominate the coculture (Fig. 2C,D, Movie S10). Thus, even using the same bacterial strains in the same media, one species can dominate in minimal flow while a different species dominates in high flow. These studies suggest that flow-dependent colonization may provide an adaptive benefit for *S. aureus*, enabling it to successfully compete with bacteria that can dominate *S. aureus* in other conditions.

**Fig. 2.**
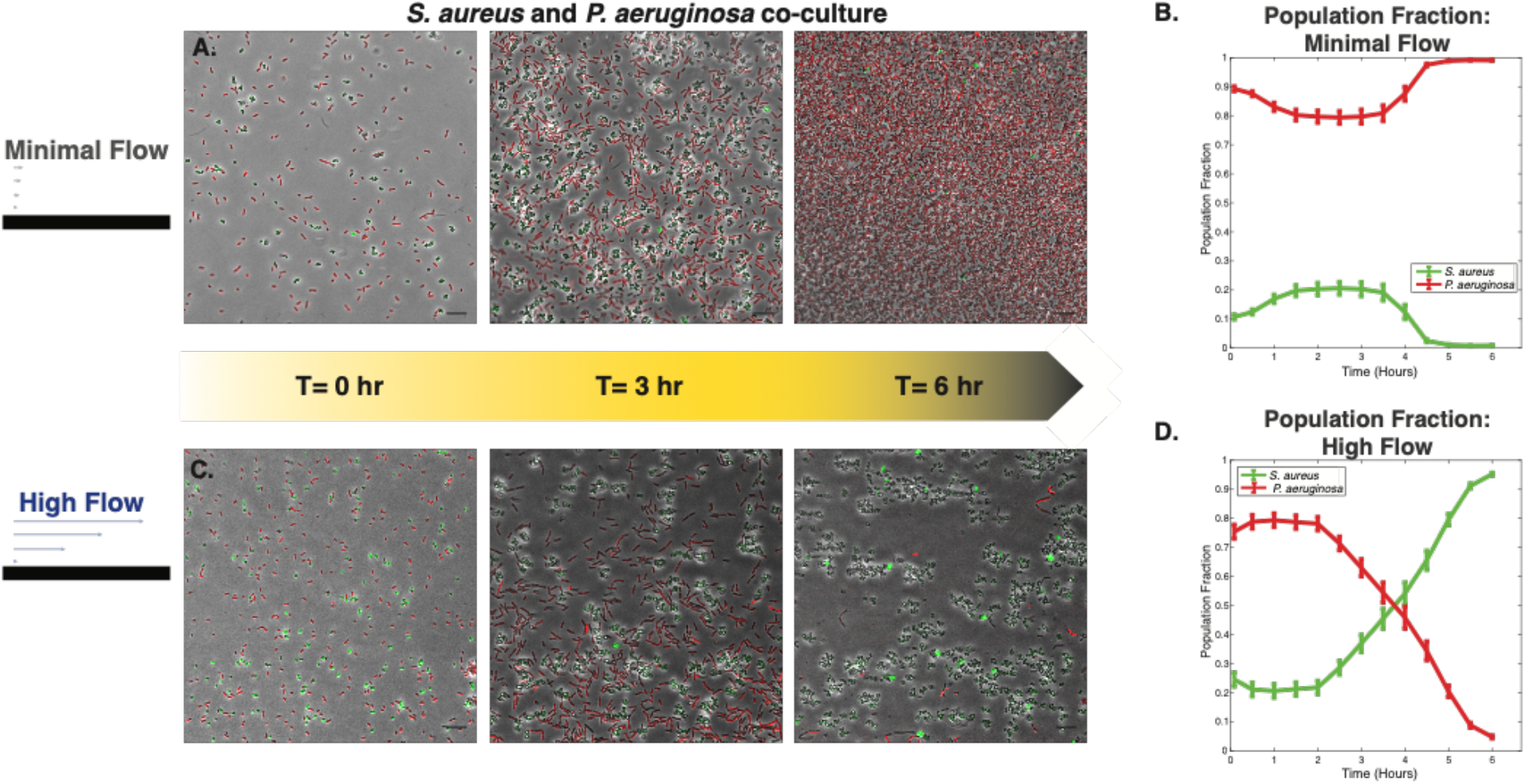
Flow-dependent colonization dynamics in co-cultures of *S. aureus* and *P. aeruginosa*. **(A)** Representative images at 0, 3 and 6 hours from a co-culture experiment of *S. aureus* (green cells) and *P. aeruginosa* (red cells) in the minimal flow condition. (B) Population fraction over the six-hour experiment run for the minimal flow condition for *S. aureus* (green) and *P. aeruginosa* (red). (C) Representative images at 0, 3 and 6 hours from a co-culture experiment of *S. aureus* (green cells) and *P. aeruginosa* (red cells) in the high flow condition. (D) Population fraction over the six-hour experiment run for the high flow condition for *S. aureus* (green) and *P. aeruginosa* (red). Scale bars for flow images are 10 μm.

### The *S. aureus* flow response is mediated by signaling molecule transport

To gain insight into the mechanisms by which flow might affect colonization, we first considered the possibility that the bacterial cells might be responding to changes in their chemical environments. Bacteria are known to produce and sense various molecules whose concentration changes can activate or repress signaling pathways that can alter the expression of factors like adhesins (*29*) that influence colonization (*24*). To test whether such chemical changes could explain the behaviors we observed for *S. aureus* and *E. faecalis*, we generated “conditioned” media where each bacterial species was grown overnight, sterile-filtered to remove intact bacterial cells, and diluted 1:1 with fresh media to ensure access to the nutrients necessary for growth. We then examined bacterial colonization in the presence of high shear rate conditions (high flow) as above, but switched the media halfway through the experiment from fresh to conditioned media. In the presence of high flow, conditioned media caused opposite effects on *S. aureus* and *E. faecalis*. Specifically, conditioned media decreased colonization for *S. aureus*, leading to colonization in high flow that more closely resembled *S. aureus* colonization in low flow (Fig. 3A, B). Meanwhile, for *E. faecalis*, conditioned media increased colonization, exacerbating the difference between high and low flow (Fig. 3C, D). Thus, the differential responses in the dynamics of *S. aureus* and *E. faecalis* colonization in flow correlate with differences in these species’ responses to conditioned media.

**Fig. 3.**
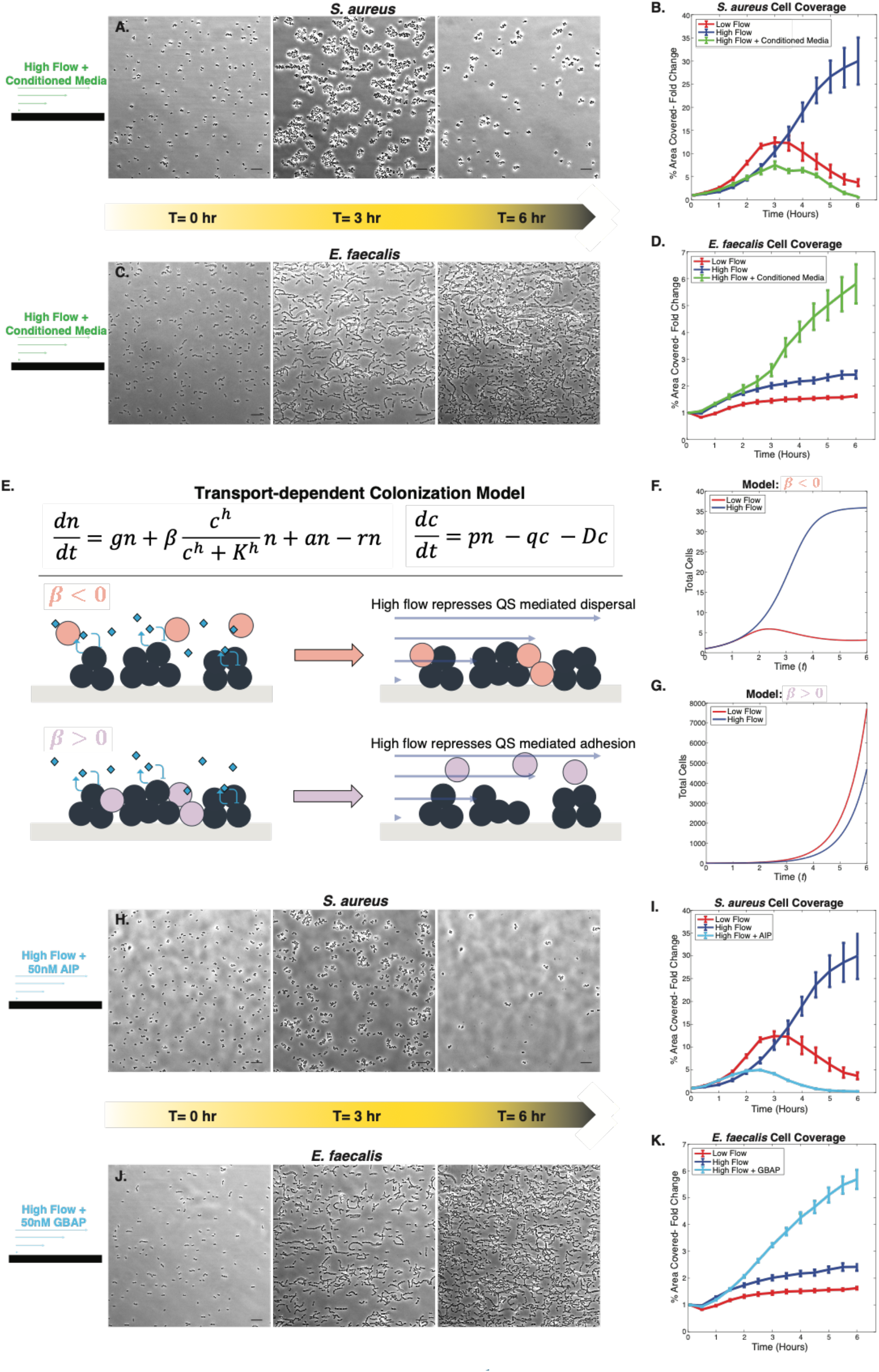
High flow transports signaling molecules *S. aureus* uses for dispersal. **(A)** Images of *S. aureus* high shear rate (400/s) experiment at 0, 3, and 6 hrs. At 3 hours, the flow was switched to conditioned media. (B) Fold changes of *S. aureus* percent area covered of high + conditioned media (green) flow experiments, with initial low (red) and high (blue) flow experiments shown for comparison. (C) Representative images of *E. faecalis* high shear rate (400/s) experiment at 0, 3, and 6 hours with the addition of conditioned media at the 3 hour timepoint. (D) Fold changes of *E. faecalis* percent are covered of high + conditioned media (green) flow experiments, with initial high (blue) and low (red) flow experiments shown for comparison. (E) Equations for the transport-dependent colonization model and schematic of the effects of high flow transport of signaling molecules. Depending on the sign of the *β* term in our model, the response under flow differs. (F) Model prediction of dispersal in low flow with a negative *β* term. (G) Model prediction of increased attachment in low flow with a positive *β* term. (H) Images of *S. aureus* at 0, 3, and 6 hours of high flow (shear rate 400/s) with media supplemented with 50 nM AIP for the entire experiment. (I) Fold change of *S. aureus* percent area covered of high + 50 nM AIP (light blue) flow experiments, with initial low (red) and high (blue) flow experiments shown for comparison. (J) Representative images of *E. faecalis* high shear rate (400/s) experiment at 0, 3, and 6 hours with media supplemented with 50 nM GBAP for the entire experiment. (K) Fold change of *E. faecalis* percent area covered of high + 50 nM GBAP (light blue) flow experiments, with initial low flow (red) and high flow (blue) experiments shown for comparison. Image scale bars are 10 μm. Error bars are SEM.

One way in which bacteria are known to respond to conditioned media is through quorum sensing (QS). In QS, bacteria both secrete and sense signaling molecules called autoinducers. In well-mixed environments, autoinducer concentration correlates with bacterial cell density, thereby enabling bacteria to drive collective behaviors. Flow can influence QS and other signaling systems by affecting small molecule transport, here defined as the movement of signaling molecule into or out of the bacterial biofilm by diffusion or advection by flow (*30*–*33*). To determine whether the interactions between flow and QS could explain the flow-dependent surface colonization of *S. aureus* and *E. faecalis*, we developed a model in which the number of cells on a surface changes as a function of cell growth, attachment and detachment, and autoinducer concentration changes as a function of the number of cells, transport, and diffusion (Fig. 3E). In this model autoinducer concentration can positively or negatively influence attachment (the parameter *β*), but autoinducer concentration does not affect growth because we observed no changes in growth rate in our experiments (Fig. 1B).

Exploring the parameters of our transport-dependent colonization model revealed that whether quorum sensing positively or negatively influences attachment (the sign of *β*) was sufficient to explain some, but not all, of the dynamics we observed experimentally. In all cases the system starts with low cell numbers, such that the effect of quorum sensing becomes more pronounced at later times. Regardless of the sign of *β*, increased shear rate led to more autoinducer transport (represented by the transport term, *q*) and thus less autoinducer accumulation. When *β* is negative and shear rate is low, growth rate initially dominates so cell numbers increase at early timepoints, but after cell density increases, autoinducer accumulates and inhibits cell attachment to an extent that eventually overwhelms growth and decreases cell numbers. In contrast, when *β* is negative and shear rate is high, autoinducer accumulates less and never overtakes growth, such that cell number steadily increases throughout the experiment (Fig. 3F). These results suggest that the effects of shear rate on QS can explain the behaviors we observe for *S. aureus* in which colonization initially increases in both high and low flow but then later diverges, continuing to increase in high flow but decreasing in low flow.

We also sought to determine if our transport-dependent colonization model can explain the colonization behavior of *E. faecalis*. Since *E. faecalis* colonization is not seen to decrease over time in any conditions, the negative *β* model that works for *S. aureus* does not work for *E. faecalis*. Because QS is known to stimulate *E. faecalis* adhesion (*34*) and we observed that conditioned media enhanced colonization (Fig. 3C-D), we examined colonization dynamics in our model when *β* is positive. In this case, we found that cell number always increased over time but did so more rapidly in low flow than in high flow because autoinducer accumulates more and thus more strongly promotes attachment in low flow (Fig. 3G). In contrast, our experimental results demonstrated that *E. faecalis* cells colonized more in high flow than in low flow (Fig. 3F-H). These results suggest that the transport-dependent colonization model cannot explain the colonization behavior of *E. faecalis* in flow.

To directly test our predictions that QS can explain the effects of flow on the colonization of *S. aureus* but not *E. faecalis*, we synthesized the primary autoinducer signaling molecule from each species: autoinducing peptide (AIP), which activates *agr* quorum sensing in *S. aureus*, and gelatinase biosynthesis-activating pheromone (GBAP), which activates *fsr* quorum sensing in *E. faecalis*. Addition of AIP to *S. aureus* decreased colonization and caused cell coverage in high flow to more closely resemble that in low flow (Fig. 3H, I). These results are consistent with previous reports that AIP signaling downregulates adhesin expression in *S. aureus* (*35*–*37*). Furthermore, genetically disrupting AIP secretion with an *agrB* mutant (*38, 39*) eliminated *S. aureus* dispersal in low flow (Fig. S2). By contrast, GBAP addition to *E. faecalis* under flow conditions increased colonization, expanding the colonization difference between the low and high flow conditions (Fig. 3J, K). Together, our findings indicate that the effects of flow on AIP QS explain the paradoxical colonization behavior of *S. aureus* but that a different mechanism must be present in *E. faecalis*.

### Mechanics of chained cells in flow drive *E. faecalis* colonization dynamics

How could high flow enhance *E. faecalis* colonization independently of QS? One striking difference between *S. aureus* and *E. faecalis* is that they have distinct growth patterns that result in different microcolony morphologies. *S. aureus* cells divide in alternating division planes (*40*), resulting in rounded microcolonies of closely clustered cells. By contrast, *E. faecalis* cells divide along a single plane (*41*), resulting in linear microcolonies of cell chains. The mechanical effects of flow on this chained microcolony morphology could conceivably help to explain the flow dependence of *E. faecalis*. Specifically, in the presence of flow, a cell chain that is anchored to the surface at one end will experience a torque produced by the shear force from the flow. The higher the flow rate, the larger this shear force becomes, increasing the torque and pushing the chain closer to the surface.

We thus sought to test whether *E. faecalis* colonization depends on shear force. Shear force can be increased independently of shear rate by increasing the viscosity of the fluid, so we increased the viscosity of the bacterial media roughly 5-fold by supplementation with 10% ficoll, a polymer that does not affect bacterial growth. This increased viscosity significantly enhanced *E. faecalis* colonization at equivalent shear rates, indicating that *E. faecalis* colonization is indeed dependent on shear force (Fig. 4 A-B). We also examined whether increased shear force pushes *E. faecalis* chains more towards the surface (as illustrated in Fig. 4C). We observed that cell chains in high shear force conditions were more sharply in focus in our images, indicating that they were closer to the focal plane positioned at the surface (Fig. 4D). To quantify this effect, we measured the fraction of total cells in focus. Relative to low flow, increasing shear force by either increasing shear rate or viscosity resulted in a significant increase in the fraction of cells in focus (Fig. 4E). We also noted that the fraction of cells in focus increased over time during each experiment (Fig. 4E). This result is consistent with our hypothesis, as cell chains elongate during the experiment and longer chains experience higher torque.

**Fig. 4.**
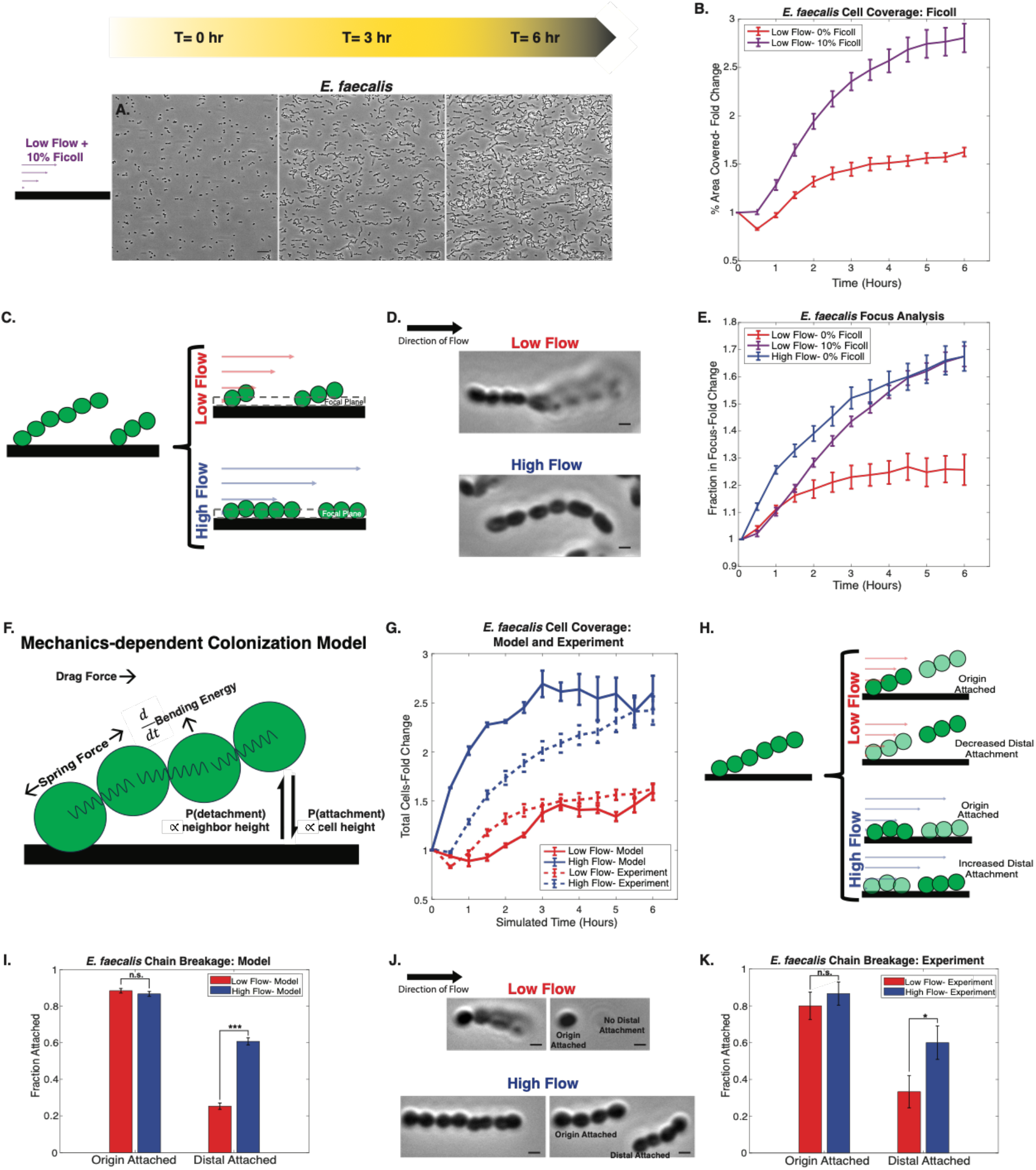
Mechanical dynamics of chains in flow lead to more cells attached and in focus in *E. faecalis*. **(A)** Representative images of *E. faecalis* low shear rate (40/s) experiment at 0, 3, and 6 hours with 10% ficoll added to the media. (B) Fold changes of *E. faecalis* percent area covered of low flow experiments with 10% ficoll (purple). Low flow with 0% ficoll (red) is shown for comparison. For flow experiment images, scale bars are 10 μm. (C) Schematic demonstrating chain dynamics and focus plane for low versus high flow. (D) Example chains in low flow (top) and high flow (bottom). In high flow, the chain is pushed to the surface and more individual cells are in focus. (E) Fold change of fraction in focus for *E. faecalis* experiments of low flow with 0% ficoll (red), 10% ficoll (purple), and high flow with 0% ficoll (blue). Brightfield images were thresholded to determine fraction in focus. (F) Schematic of the forces and attachment/detachment dynamics implemented in our mechanics-dependent colonization model. (G) Comparison of total cell coverage from the model in low and high flow (red and blue, solid line) compared to experimental *E. faecalis* runs (red and blue, dashed line). (H) Schematic depicting possible chain dynamics following a breakage event in low (upper two) and high (lower two) flow. (I) Mechanics model prediction of chain breakage dynamics in low flow conditions (red) and high flow conditions (blue). Following a breakage event in the model, the anchor and distal attachments were recorded for 100 simulated chains. **(J)** Experimental observations of breakage events from low flow (top), where the chain breaks and only the origin cell is still attached, and high flow (bottom), where the chain breaks and both the distal and origin cells are attached. Images shown are from two consecutive time points (Δ*t* = 5 min). (K) Fraction attached for both origin and distal in high and low flow experiments. Flow videos were analyzed for breakage events throughout the 6 hour experiment. For *n* = 30 chains in both high and low flow, the outcome after a breakage - anchor or distal adherence - was counted. We note that chains could break and have both origin and distal adhered or could completely detach from the surface, leading to neither origin nor distal adhered. P value was determined using a paired t-test. * = *p* <0.05, *** = *p* < 0.005. Scale bars in both D and J are 1 μm. For all graphs, error bars are SEM.

To mechanistically explore the mechanical responses of *E. faecalis* chains to flow, we developed a mechanics-dependent colonization model (Fig. 4F). This is a 2D agent-based model in which cells are agents that can be connected to one another to form chains. Chains are allowed to grow, as growth continues during our experiments, and each cell can independently attach or detach from the surface. The attachment probability depends on a cell’s distance from the surface, while detachment probability increases if neighboring cells are detached. Moreover, the connection between two neighboring cells in a chain can stochastically break. A chain is considered to be attached to the surface if any of its cells are attached. To mimic the dynamics observed experimentally, chains begin with an origin cell attached to the surface and subsequent cells are attached at the end of the chain by a stiff spring with a bending energy that keeps the cells in a relatively straight line. An additional force acting on the chains is a drag force that acts in the horizontal direction to recapitulate the shear force from flow. We found that in our simulations, increased flow rate increased the number of cells that attached to the surface (Fig. 4G), recapitulating our experimental results with *E. faecalis*. Our simulations also revealed that the effect of flow on *E. faecalis* colonization could be explained by the impact of flow on the outcomes of chain breakage events. When a chain breaks, the part of the chain proximal to the ancestral origin cell has the same probability of remaining attached regardless of the flow rate (Fig. 4I). This portion of the chain has a high probability of being attached in all cases (Fig. 4H). In contrast, the part of the chain distal to the origin cell has a higher probability of remaining attached in high flow than in low flow (Fig. 4I). This results from the higher drag force pushing the cell chain closer to the surface and thus increasing the probability that cells in the distal part of the chain will be attached to the surface (Fig. 4H).

To validate the mechanics-dependent colonization model’s prediction that chain breakages should have flow-dependent outcomes, we analyzed chain breakage events from our experiments in both high and low flow. Specifically, we quantified the fraction of breakages where the origin-proximal cells or distal cells remained adhered after the break (Fig. 4J). As predicted, there was no significant difference between the fraction of origin cells adhered following a breakage between low and high flow conditions (Fig. 4K). However, there was a significant difference in distal cell adherence, with more distal cells adhering in high flow than in low flow after a breakage event (Fig. 4K). Examples of the differences in chain breakage outcomes in high and low flow are shown in Movies S11-14.

Our mechanical model also predicted that strains that formed longer chains should increase colonization in both low and high flow (Fig S3C). To experimentally test this prediction we examined colonization dynamics in an *E. faecalis ΔatlA* mutant (*42*), that has longer cell chains due to the lack of the peptidoglycan hydrolase AtlA. The *ΔatlA* mutant displayed increased surface colonization in both low and high flow conditions (Fig S3A, B, D). *ΔatlA* mutant cells increased distal attachment following breakages in low flow (Fig S3E) but showed no significance difference between the origin-proximal cells attached after a breakage (Fig S3E). Together, these results establish the mechanism of *E. faecalis* flow-dependent colonization: high flow rates mechanically push *E. faecalis* chains towards the surface, resulting in increased colonization by increasing attachment and thereby reducing cell dispersal upon chain breakage.

## Discussion

Fluid flow introduces complexities to bacterial dynamics that can result in unexpected emergent behaviors. Here we elucidate two different mechanisms by which two species preferentially colonize surfaces in higher flow environments. Previous studies of bacterial colonization in flow have primarily focused on the mechanisms by which single cells adhere to surfaces, such as the catch bonds formed between Uropathogenic *Escherichia coli* and the mannosylated surface of epithelial cells (*22*) or the mechanosensitive Type IV pili of *Pseudomonas aeruginosa* (*43, 44*). By contrast, here we find that increased flow stimulates colonization of both *S. aureus* and *E. faecalis* via distinct mechanisms that each function at the multicellular scale. For *S. aureus*, but not *E. faecalis*, flow enhances colonization through a reduction in QS signaling resulting from increased autoinducer transport, which in turn leads to adhesin downregulation and cell detachment (*32, 35, 45*). Importantly, whereas previous studies on quorum sensing in flow suggested that flow negates the physiological impact of quorum sensing in many contexts, our findings suggest that bacteria could harness the inhibition of quorum sensing in flow as a beneficial adaptation to promote colonization. The potential adaptive benefit of flow-dependent colonization is supported by competition experiments between *S. aureus* and *P. aeruginosa*, another pathogen that is often found with *S. aureus* but does not commonly cause endocarditis. Specifically, whereas *P. aeruginosa* outcompetes *S. aureus* in minimal flow, *S. aureus* outcompetes *P. aeruginosa* in high flow. Additional studies inspired by this work will help to determine the importance of interactions between multicellular signaling and flow in more complex environments like heart valves.

In contrast to *S. aureus*, we found that mechanical effects on chained-cell microcolonies were sufficient to explain how flow stimulates the colonization of *E. faecalis*. Altering fluid viscosity indicated that the *E. faecalis* colonization effect depends upon shear force rather than solely the shear rate, and agreement between predictions generated by a biophysical mechanics-dependent colonization model and experimental observations confirmed this hypothesis. Specifically, we found that the linear chains of cells produced by *E. faecalis* microcolonies experience a torque that is proportional to viscous effects. Consequently, higher flow pushes the cells more towards the surface, leading to increased attachment, and a mutant that increases chain length enhances colonization.

Interestingly, while *S. aureus* and *E. faecalis* achieve flow-dependent colonization in different ways, both mechanisms do not operate on single cells but rather depend on the specific collective morphologies of each species’ microcolonies. The clustered multicellular morphology in which *S. aureus* cells grow facilitates the local 3D signaling interactions that mediate quorum sensing. Meanwhile, the linear multicellular morphology of *E. faecalis* produces a lever-like effect that promotes the torque required for flow-dependent attachment. Many studies of single-cell bacterial morphology have noted the importance of individual cell shape (*46*–*48*), but our findings suggest that even when individual bacteria have cells of the same shape, there can be adaptive benefits to the collective morphologies that form in their multicellular microcolonies.

## Acknowledgements

We thank members of the Gitai Lab, Josh Shaevitz, and members of the Shaevitz Lab for discussion and feedback.

## Author Contributions

Conceptualization: KMH, ZG

Methodology: KMH, ZG

Chemical Synthesis: SPB, TWM

Investigation and Analysis: KMH

Computational Modeling: KMH, HAS, NSW

Funding acquisition: HAS, ZG

Supervision: ZG

Writing – original draft: KMH, ZG

Writing – review & editing: KMH, SPB, HAS, TWM, NSW, ZG

## Data and Materials availability

### Data Availability

The data supporting the findings of the study are available in this article and its Supplementary Information. Additionally, the raw data that support the findings of this study are available from the corresponding author upon request.

### Code Availability

Custom Fiji Macros and MATLAB code used for cell quantification and focus analysis of the microscopy data are freely available from the corresponding author upon request. Similarly, MATLAB code for both the transport-dependent colonization model and mechanics-dependent colonization model are freely available from the corresponding author upon request.

## Supplementary Materials

Materials and Methods

Supplementary Text

Figs. S1 to S5

Tables S1 to S3

References (#*49-53*)

Movies S1 to S15

