## Supplemental Information, Tables, and Figures for "Distinct microcolony morphologies promote flow-dependent bacterial colonization"

**The PDF file includes:**

Materials and Methods
Supplementary Text
Figs. S1 to S5
Tables S1 to S3
References 49-53

**Other Supplementary Materials for this manuscript include the following:**

Movies S1 to S15

### Materials and Methods

#### Bacterial Strains and Growth Conditions

*S. aureus* experiments used the strain USA300 MRSA. The *S. aureus agrB* transposon mutant was from the Nebraska Transposon Mutant Library (*38*, *39*). Constitutive green fluorescent *S. aureus* USA300 used for co-culture experiments was from (*49*)(generously provided by the Torres Lab at NYU). *E. faecalis* studies were done using the strain OG1RF, fluorescently labeled with the plasmid pBSU101-GFP (*50*) (generously provided by the Wood Lab at the University of Michigan). The *E. faecalis ΔatlA* mutant used was from (*42*). Constitutive red fluorescent *P. aeruginosa* PA01 was from (*43*). Wild type species used for supplemental studies in low and high flow were *E. coli* MG1655 and *S. pneumoniae* TIG4 (generously provided by the Veening Lab at the University of Lausanne)

Media was prepared according to manufacturer recommendation. Overnight cultures were grown in floor shakers at 37C in Luria-Bertani Miller (LB) Broth (BD Biosciences) for *S. aureus*, *P. aeruginosa*, and *E. coli* or Brain Heart Infusion (BHI) Broth (BD Biosciences) for *E. faecalis* and *S. pneumoniae* following isolation of a single colony from LB or BHI agar plates, respectively. For *E. faecalis* OG1RF, the plasmid pBSU101-GFP was maintained with the addition of 120 μg/ml of the antibiotic Spectinomycin (Sigma) in the plates as well as growth media.

Following overnight growth, cells were diluted in fresh media to an optical density (OD) between 0.40 and 0.45 and loaded into plastic syringes with a 27G needle in preparation for microfluidic chip experiments. For mixed co-culture experiments, both *S. aureus* and *P. aeruginosa* were diluted in fresh media to an OD between 0.40 and 0.45 before mixing together 1:1. This mixture was then loaded into plastic syringes with a 27G needle in preparation for microfluidic chip experiments.

For conditioned media experiments, overnight cultures were centrifuged at 9000xg for 5 minutes to pellet the cells. The supernatant was removed and filtered using 0.2 μm filter disks (Pall Corporation) to ensure all cells had been removed. This filter sterilized overnight media was then mixed 1:1 with fresh LB (*S. aureus*) or BHI (*E. faecalis*) to create our conditioned media. For conditioned media experiments, we switched the syringes to conditioned media syringes from fresh media following 3 hours of the flow run.

For *S. aureus* AIP experiments, synthesized AIP-I (obtained as described below) was added to fresh LB to a final concentration of 50 nM. The LB supplemented with AIP was used for the entire flow experiment. Similarly, for *E. faecalis* GBAP runs, synthesized GBAP was added to fresh BHI to a final concentration of 50 nM. Again, the BHI supplemented with GBAP was used for the entire 6-hour flow experiment.

For *E. faecalis* viscosity and shear force experiments, 10% ficoll (Sigma) was added to BHI. This BHI + 10% ficoll media was used for the entire 6-hour experiment.

#### AIP and GBAP Synthesis

*AIP:* AIP-I was synthesized via a protocol adapted from Zhao *et al.* 2022 (*51*). Briefly, we performed standard automated Fmoc-SPPS on a Liberty Blue microwave-assisted peptide synthesizer (CEM), utilizing a DIC-Oxyma coupling strategy on a hydrazine derivatized Cl-TCP(Cl)-ProTide resin (CEM). After SPPS, cleavage of the resin yielded a linear peptide with a C-terminal hydrazide. This hydrazide was oxidized with 15 eq. of $NaNO_2$ in a 6 M guanidinium hydrochloride buffer solution at pH 3. A peptide thioester was formed upon the addition of 15 eq. MESNa at pH 6.2. The MESNa thioester was purified by RP-HPLC and lyophilized. To generate the cyclized final product, the peptide thioester was solubilized in PBS with 5 mM TCEP at pH 7. AIP-I was purified again by RP-HPLC, lyophilized, and taken up in DMSO. Concentration was determined by NMR relative to known standards. (Fig. S4)

*GBAP*: GBAP was synthesized via a protocol adapted from McBrayer *et al.* 2017 (*52*). Briefly, GBAP was generated by Fmoc-SPPS using a HATU coupling strategy extending the chain from Gln9 of the peptide. The unprotected side chain of Fmoc-Glu-Oall was coupled to a H-Rink amide ChemMatrix® resin (Matrix Innovations). The peptide was extended to Asn2 by standard Fmoc-SPPS. To the N-terminus was coupled Boc-Gln-OH to supply Gln1 and cap this chain of the peptide. To add Met9 and Trp10 and form the ester linkage of the peptide lactone, the O-trityl protecting group of Ser3 was first removed with 1% TFA. Fmoc-Met-OH (10 eq) was coupled to the Ser3 side chain by a Steglich-type esterification, utilizing DIC (9.9 eq), DIEA (10 eq), and DMAP (0.5 eq). After validating the formation of the Met9 ester by LC-MS, Trp10 was added by standard Fmoc-SPPS. The C-terminal Oalloc protecting group on Gln9 was removed by treatment with 10 eq 1,3-dimethybarbituric acid and 0.3 eq tetrakis(triphenylphosphine)palladium(0) in DCM. After repeating the Alloc deprotection, the Pd catalyst was removed by washing with 1% sodium diethylthiocarbamate. On-resin cyclization was performed with PyOxim (1.25 eq) and DIEA (2.5 eq). Cyclized product was cleaved from the resin and purified by RP-HPLC. Pure products were lyophilized, taken up in DMSO, and quantified by NMR. (Fig. S4)

**Microfluidic Devices**

Flow channels were designed in Blender and a 3D mold was printed by Protolabs (Maple Plain, MN). Each chip had six channels that were 2 cm long x 500 μm wide x 100 μm tall with separate inlet and outlet ports for each channel. Briefly, Polydimethylsiloxane (PDMS) was poured onto the 3D mold, baked overnight, and then plasma-bonded to 22x44 mm #1.5 glass coverslips (Avantor). Following this, the completed chips were baked overnight at 65C to complete the bonding before use.

**Flow Experiments**

Microfluidic chips were prepared for flow experiments by adding inlet and outlet tubing (BD Intramedic Polyethylene Tubing .015” ID 0.043” OD) and loading the channel with media before addition of our diluted cells. Following cell addition, a plastic media-filled syringe with 27G needle was connected to the inlet tubing and mounted on a syringe pump (New Era Pump Systems, Inc.) and the outlet tubing was placed into a waste container to collect the outflow. Following 15-20 minutes of initial attachment time, flow was turned on at one of two rates. For high flow conditions corresponding to a shear rate 400/s, a flow rate of 20 μl/min was used. For low flow conditions corresponding to a shear rate 40/s, a flow rate of 2 μl/min was used.

For *S. aureus* and *P. aeruginosa* co-culture experiments, the high flow conditions were as described above. For minimal flow, following initial attachment of the channels, flow was turned on at a flow rate of 35 μl/min for five minutes to clear the channels of excess, unattached cells. For minimal flow experimental runs, flow rate was lowered to 0.1 μl/min, a shear rate of 2/s, to keep the channel clear and provide fresh media throughout the run.

**Phase Contrast and Fluorescence Microscopy**

For all flow experiments, a Nikon 90i inverted microscope equipped with a 100x 1.4 N.A. objective and with an incubator held at 37C was used. Microscope control and image acquisition was done using NIS Elements (Nikon, version 4.60.00). Five positions spread along each channel were chosen and the image plane was focused on the glass coverslip. Flow was turned on following the first set of images. Images were then taken every 5 minutes for a total of 6 hours for both species. Phase contrast images were taken for all species. For *E. faecalis*, fluorescent images were taken using a GFP filter. To obtain doubling time information (Fig 1B), the same flow set up and microscopy filters were used, but images were taken every 2 minutes for better doubling time resolution.

**Quantification of Cell Coverage**

Cell coverage analysis was completed using a custom image analysis macro in Fiji and MATLAB code (software available upon request, Mathworks, Natick, MA). Briefly, for *S. aureus*, cell edges were found and contrast between the edges and background was enhanced using a CLAHE filter. Images were thresholded to remove background, binarized, and edges were filled in. Total cell coverage was determined by summing the total pixel count of the binarized images. For *E. faecalis*, analysis was done using fluorescent images to better capture the chains. Images were run through the CLAHE filter before having the background subtracted and binarized. Once again, total cell coverage was then determined by summing the total pixel count of the binarized images. The measured cell coverages were then loaded into MATLAB. The total coverage was normalized to the first image for each experiment position, and then the normalized total coverage for all positions of each experimental condition were averaged together and plotted over the experimental time. Error bars on these averages are reported as standard error on the mean (SEM).

Population fractions of the mixed co-cultures of *S. aureus* and *P. aeruginosa* were again analyzed using custom image analysis pipelines using Fiji and MATLAB. Briefly, fluorescent images for each species were background subtracted, enhanced using a CLAHE filter and binarized. Cell coverage for each species was determined by summing total pixel count and these measurements were loaded into MATLAB. Population fraction was found by dividing the individual species coverage by the total coverage of both species at each time point. The population fractions from multiple experimental runs were then averaged together for each experimental condition (minimal or high flow). Error bars are reported as SEM.

**Focus Analysis**

*E. faecalis* focus analysis was again completed with a custom pipeline in Fiji and MATLAB (software available upon request). Analysis began in Fiji, with the phase contrast images CLAHE filtered followed by finding the edges of the cells. These edged images were then loaded into MATLAB. The edges of out-of-focus cells were dim and could be excluded by thresholding the images. The same threshold was used for all conditions. For each time point in an experiment, the thresholded and non-thresholded images were binarized and the total pixel value of each binarized image was calculated. The fraction in focus was then calculated by dividing the thresholded image total by the non thresholded image. These results were then normalized to the initial time point for each position and all positions for a specific condition were averaged together. Error bars are again reported as SEM.

**Chain Breakage Analysis**

For analysis of chain breakage outcomes, thirty breakage events were calculated by hand for low and high flow. Following a breakage, origin attachment and/or distal attachment was recorded. The fraction of events where origin or distal attachment was recorded was calculated by dividing the total number of times the breakage outcome was recorded by our total number of breakage observations (thirty). Error bars are SEM. Significance was determined used a paired t-test.

For simulations, anytime a breakage event was occurred, the outcome (origin and/or distal attachment) was recorded. Following the end of a chain simulation, these outcomes were recorded. For 100 chains in each simulation set, representing a 'position' along a simulated flow channel, the total outcomes of each were calculated and again the fractions were calculated, error bars are SEM, and significance was determined using a paired t-test.

**Supplementary Text**

***S. aureus* transport effect captured in a model of ODEs**

To incorporate the dynamics of QS systems and transport of signaling molecules, we developed a model of coupled ODEs, describing cell number and growth (Eq. 1) and production and transport of the signaling molecule (Eq. 2). For the number of cells $n$, we use

$$\frac{dn}{dt} = gn(t) + \beta\frac{c^h}{c^h + K^h}n(t) + an(t) - rn(t) \quad \text{Eq. 1}$$

where, $t$ is time, $g$ is the cellular growth rate, $\beta$ is the dispersal constant, $c$ is the concentration of the autoinducer signaling molecule, $a$ is the rate of attachment of new cells, and $r$ is the quorum sensing-independent detachment of cells. We note $a$ and $r$ experimentally appear to be negligible (see Movie S15: dynamics of *S. aureus* in the absence of growth) and were thus set to 0 for our model simulations. We use a Hill function to describe the dynamics of the QS system, as a function of the signaling molecule concentration. Here, $K$ is the half maximal concentration and $h$ is the Hill coefficient. As the shape of the QS response is system specific, we modeled a range of $h$ values with both positive and negative $\beta$ (Fig. S5). We note the overall behavior is qualitatively insensitive to the precise value of $h$. In the main text, we use $h = 3$ to describe the positive feedback that is characteristic of QS systems. Also, for autoinducer concentration $c$ we use

$$\frac{dc}{dt} = pn(t) - qc(t) - Dc(t) \quad \text{Eq. 2}$$

where $p$ as the production rate of the molecule, $q$ is the transport rate of the molecule (analogous to our flow rate), and $D$ represents the diffusion away from the colony. For our simulations, we assume $D << q$ and set $D$ to 0. The parameters used for simulations of this model can be found in Table S1.

**Model describing forces on a chain mimics *E. faecalis* flow dynamics**

To probe the mechanical effects of flow on *E. faecalis* cell chains, we created a computational model of *E. faecalis* chains in flow modeling a spring force between the cells, a bending energy keeping the cells in line, and a drag force accounting for flow. To determine the different responses in low and high flow, we ran simulations where the only difference in conditions was the drag force: this force was 10x larger in high flow simulations, to account for the 10x larger flow rate in high flow experiments.

Our agent-based model describing the mechanics-dependent colonization of *E. faecalis* was written in MATLAB. Each simulated chain was initialized to a random length between one and five cells long, with cells, considered simply as vertices in the chain, connected to their neighbors by a stiff harmonic spring with spring constant $k_s$. For chains of two cells or more, the subsequent cells were initialized at an angle 45° from the surface, with this angle allowed to fluctuate during the simulations. The chain length also "grows" by the addition of one cell at the end of the chain, initialized in line with the existing chain, after a set number of timesteps. Following initialization, at each timestep the cells are moved based on the forces acting upon them. There are three forces we model: the spring force between each pair of cells acting in both the $x$ and $y$ direction, a drag force which is proportional to the shear rate and the height of the cell above the surface acting in the $x$ direction, and a derivative of the bending energy that tends to keep the cells in a roughly straight line, again acting in both the $x$ and $y$ directions. Based on these calculated forces, the $x$ and $y$ positions of the cells are updated, followed by checking for surface attachment/detachment and chain breakages. For attachment, the probability of attachment of each cell increases linearly as it gets closer to the surface. If a cell is already attached, at each step it is checked for detachment, with the probability of detaching increasing as a function of the total height of the two neighbor cells to the attached cell. Detachment of the origin cell in a chain was seen rarely experimentally and this detachment event was modeled as a stochastic process. If the origin cell detaches and no other cells in the chain are attached, the simulation is stopped and the chain is marked as both origin and distal detached. To model breakages, at each step the tension of the spring between each pair of cells is calculated. If the spring tension is larger than a set value, it has a breakage probability that increases exponentially with increasing spring length. Following a breakage along the chain, cells on each side of the breakage are checked for attachment. If any cells in the segment of the origin cell are attached, "origin segment attachment" is recorded. Similarly, if any cells on the distal chain segment are attached, "distal segment attachment" is recorded.

At each time step, the cell positions were updated according to the deterministic forces (detailed below) plus a small stochastic force in both *x* and *y* directions, sampled from Gaussian distributions centered around 0 with a standard deviation of 0.1. We note that if updating the cell positions would cause any cell to move to a negative *y* position, the cell was set to the surface plus a small positive gaussian step (from a distribution of mean 0 and standard deviation 0.1). After updating the position, each cell was checked for attachment, then detachment. If detachment occurred, the new *y* position was similarly set to a small gaussian step above the surface (from a distribution of mean 0 and standard deviation 0.01). Following these updates, chains were checked for breakages, before moving to the next time step. The results of 100 chains, simulated in either the low flow or high flow condition, were combined to mimic the corresponding flow experiment.

Regarding the spring forces on the cells in more detail, given three cells at positions *i*-1, *I,* and *i*+1 connected to each other with springs*,* the potential energy stored in the spring connecting the cell at position *I* to the cell at position *i*-1 is:

$$E_s = \frac{1}{2}k_s\left(\sqrt{\left(x_i - x_{i-1}\right)^2 - \left(y_i - y_{i-1}\right)^2} - l_0\right)^2, \qquad \text{Eq. 3}$$

where $k_s$ is the spring constant, *x* is the *x*-position of the cell at *I* or *i*-1, respectively, and *y* similarly is the *y*-position of the cell at *I* or *i*-1, and $l_0$ is the unstretched length of the spring. The energy of the spring connecting cell *I* to cell *i*+1, is given by the same expression, just replacing the *i*-1 subscripts with *i*+1. The total spring force in *x* or *y* is then found by taking the derivative of the two energy expressions with respect to $x_i$ and $y_i$ and summing the *x* or *y* expressions, respectively.

The bending energy acting on the cell at position *i* which makes an angle $\theta$ with the cells at positions *i*-1 and *i*+1 is given by the expression:

$$U_b(\theta) = k_b\left[1 + \cos(\theta)\right], \qquad \text{Eq. 4}$$

where $k_b$ is a bending constant. This expression can be rewritten in terms of the vectors $\vec{v}_{i,i-1}$ and $\vec{v}_{i,i+1}$ , where these are the vectors from position *i* to position *i*-1 and position *i* to position *i*+1, respectively:

$$U_b(\theta) = k_b\left[1 + \frac{\vec{v}_{i,i-1} \cdot \vec{v}_{i,i+1}}{\left\|\vec{v}_{i,i-1}\right\|\left\|\vec{v}_{i,i+1}\right\|}\right]. \qquad \text{Eq. 5}$$

Taking the derivative of this expression with respect to $x_i$ and $y_i$ , $x_{i-1}$ and $y_{i-1}$ , and $x_{i+1}$ and $y_{i+1}$ , gives the total bending force acting on the three cells due to the angle formed by cells *i*-1, *i*, and *i*+1.

We approximate the force from the flow condition as a drag force that acts in the *x* direction and depends on the height of the cell above the surface:

$$F_{drag,i} = d * f * y_i\,\hat{x} \qquad \text{Eq. 6}$$

where d is a constant, set to 1 for our simulations, *f* is the flow rate and $y_i$ is the *y* position of cell *i*. The parameters and probabilities used in simulations of our chains in low and high flow conditions can be found in Tables S2 and S3, respectively.

***S. aureus*** **attachment/detachment analysis**
To observe the amount of spontaneous detachment within our channels, as well as attachment from upstream detachment, we performed an experiment in which *S. aureus* was observed in flow in conditions that prevented cell growth (Movie S15). Specifically, we used a minimal growth media AAM (*53*) but without the addition of glucose that is normally added to AAM. In these experiments we observed only rare detachment and attachment events, did not see clustered morphologies, and did not see dispersal events, indicating that all of these dynamic responses rely on growth.

**Calculation of shear rate**
Microfluidic channels used in this study had a rectangular cross section. We calculated the shear rate ($\gamma$) at the floor and ceiling of the channel, where height is << width, using the equation:

$$\gamma = \frac{6Q}{wh^2}, \qquad \text{Eq. 7}$$

where *Q* is the total flow rate, *w* is the width of the channel and *h* is the channels height.

Estimates of the shear rate in a heart valve was obtained for a channel with a circular cross section. For a circular channel, the shear rate ($\gamma$) along the wall is:

$$\gamma = \frac{4Q}{\pi r^3}, \qquad \text{Eq. 8}$$

where *Q* is again the total flow rate and *r* is the radius of the cross section.

We estimated the flow rate and radius of the cross section of a valve in the heart from values given in (*26*, *27*). Focusing on values for the mitral valve, we estimate during a heartbeat that all of the fluid from the left atrium passes through the valve. The average volume of the left atrium for males is reported to be 81 ± 18 ml and for women to be 67 ± 14 ml. For our calculation of an "average" person, we estimated the volume passing though the valve to be 75 ml. We assume a resting heart rate of about 65 beats per minute, meaning the left atrium volume would pass through the valve in at most ~0.92 seconds. This gives a flow rate, *Q*, of 75 ml/0.92 s, or 81.25 $cm^3$/s. To determine the radius, it was reported that the mitral valve has a surface area of 4-6 $cm^2$. We assume the cross-sectional area of the valve is slightly larger than the cross-sectional area the fluid passes through, as the tissue making up the valve leaflets would block some of the area. We estimated a cross-sectional area of the opening for the fluid to be ~3 $cm^2$. We can calculate the radius of this area using the equation for the area of a circle, finding $r = 0.98$ cm. Putting these values of *r* and *Q* into Equation 8, we find:

$$\gamma = \frac{4Q}{\pi r^3} = \frac{4 * 81.25 \frac{cm^3}{s}}{\pi (0.98\ cm)^3} = 109.9/_s. \qquad \text{Eq. 9}$$

With this value, we roughly estimate the shear rate in valves to be around 100/s, though we do acknowledge we are assuming constant flow and a circular valve cross section.

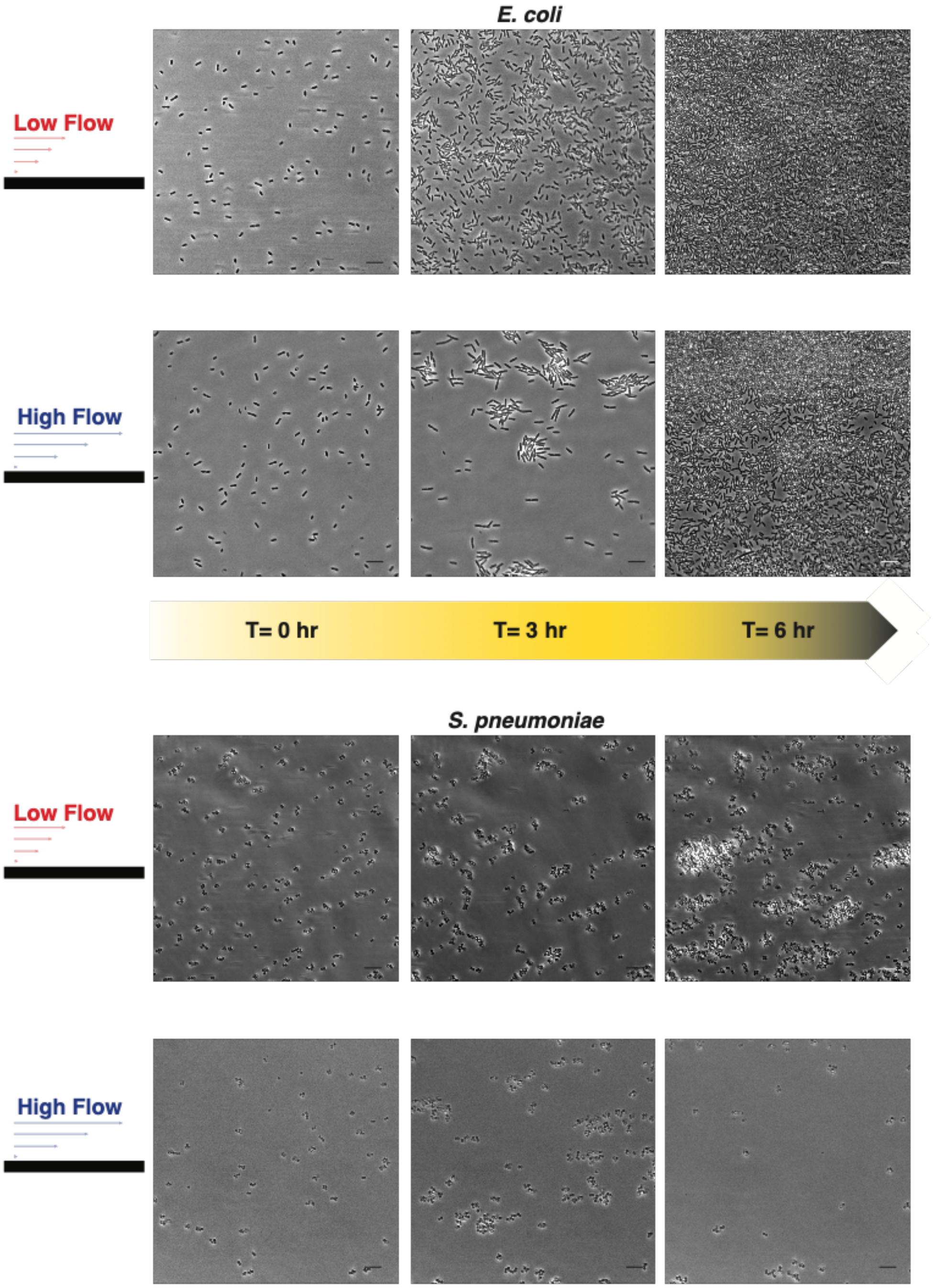


**Fig. S1. *E. coli* and *S. pneumoniae* do not follow paradoxical high flow response (A)** Representative images of a low flow *E. coli* MG1655 experiment at 0, 3, and 6 hours. (B) Representative images of a high flow *E. coli* MG1655 experiment at 0, 3, and 6 hours. (C) Representative images of a low flow *S. pneumoniae* TIG4 experiment at 0, 3, and 6 hours. (D) Representative images of a high flow *S. pneumoniae* TIG4 experiment at 0, 3, and 6 hours. Scale bars are 10 μm.

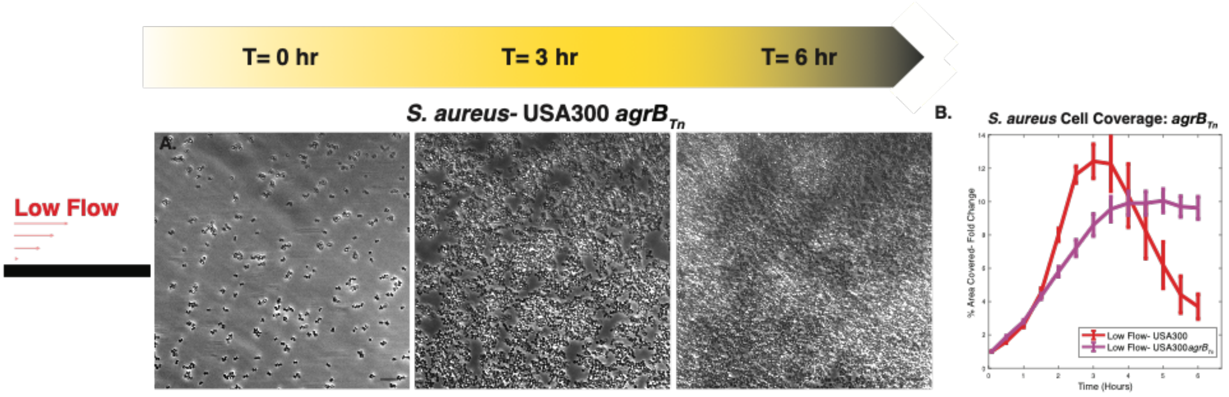


**Fig. S2.** ***S. aureus agrB*** **transposon mutant does not disperse in low flow (A)** Representative images of a low flow *S. aureus* USA300 $agrB_{Tn}$ experiment at 0, 3, and 6 hours. (B) Fold change of percent area covered of both WT *S. aureus* USA300 (red) and the mutant *S. aureus* USA300 $agrB_{Tn}$ strain (magenta). Scale bars are 10 µm.

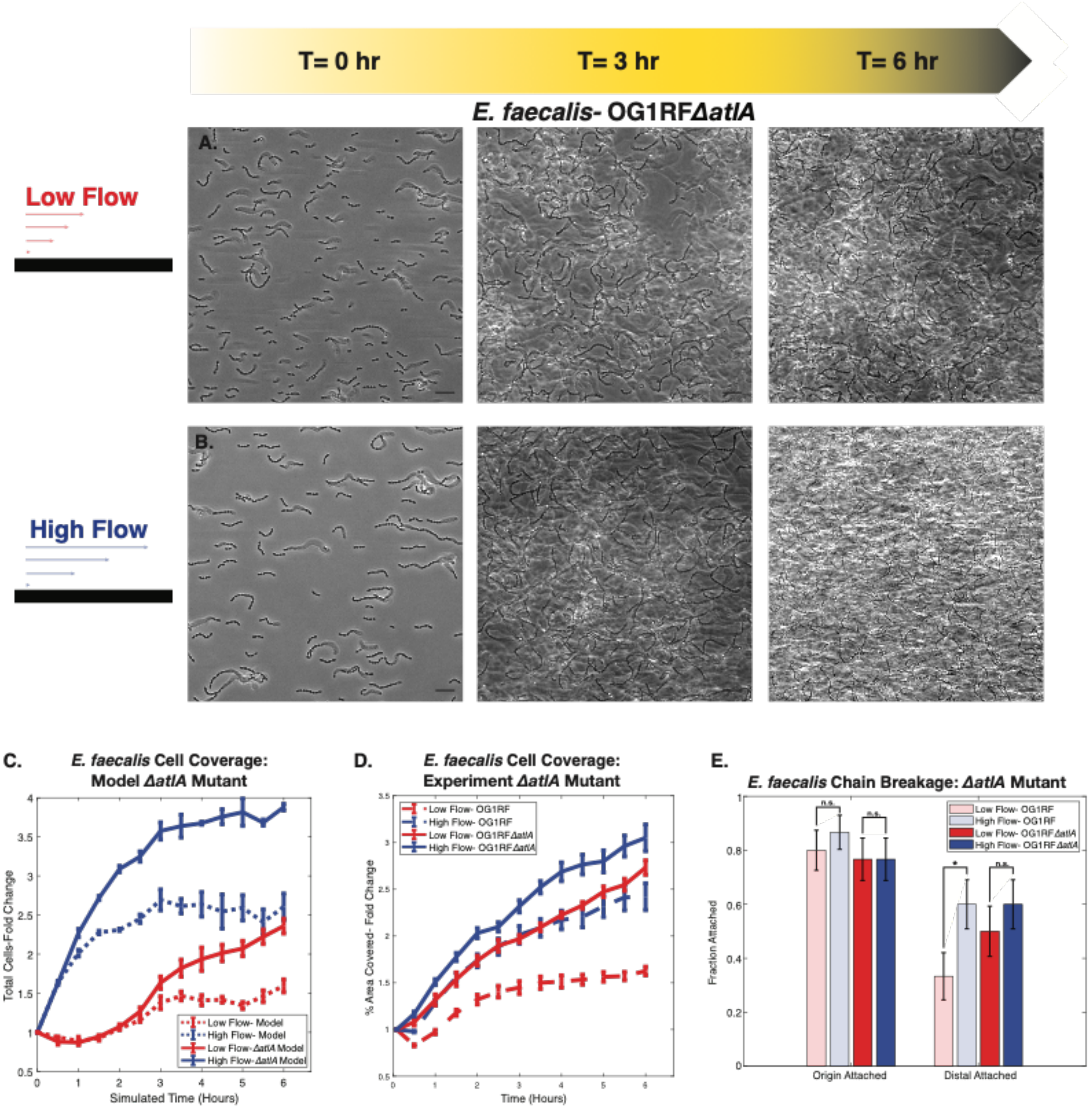


**Fig. S3. Longer chains from *E. faecalis ΔatlA* mutant leads to increased surface colonization. (A)** Representative images of a low flow *E. faecalis* OG1RF *ΔatlA* experiment at 0, 3, and 6 hours. (B) Representative images of a high flow *E. faecalis* OG1RF *ΔatlA* experiment at 0, 3, and 6 hours. (C) Fold change of total cells for our mechanics-dependent colonization model. Original model runs for low and high flow are shown in dashed lines for comparison. The breakage tension in the model was increased to explore longer chain dynamics, mimicking the *ΔatlA* mutant and leading to more cells seen in both low (red) and high (blue) flow model runs. (D) Fold change of the percent area covered of both our WT *E. faecalis* (low and high flow, red and blue dashed) and a mutant *ΔatlA* strain (low and high flow, red and blue) that alters septum cleavage leading to an even longer chained morphology. Both low and high flow for the mutant lead to a larger area covered. (E) Chain breakage analysis and comparison of WT (low and high flow, pale red and blue) to *ΔatlA* mutant (low and high flow, red and blue). Scale bars are 10 μm. Error bars are SEM. P value was determined using a paired t-test: * = $p$ <0.05.

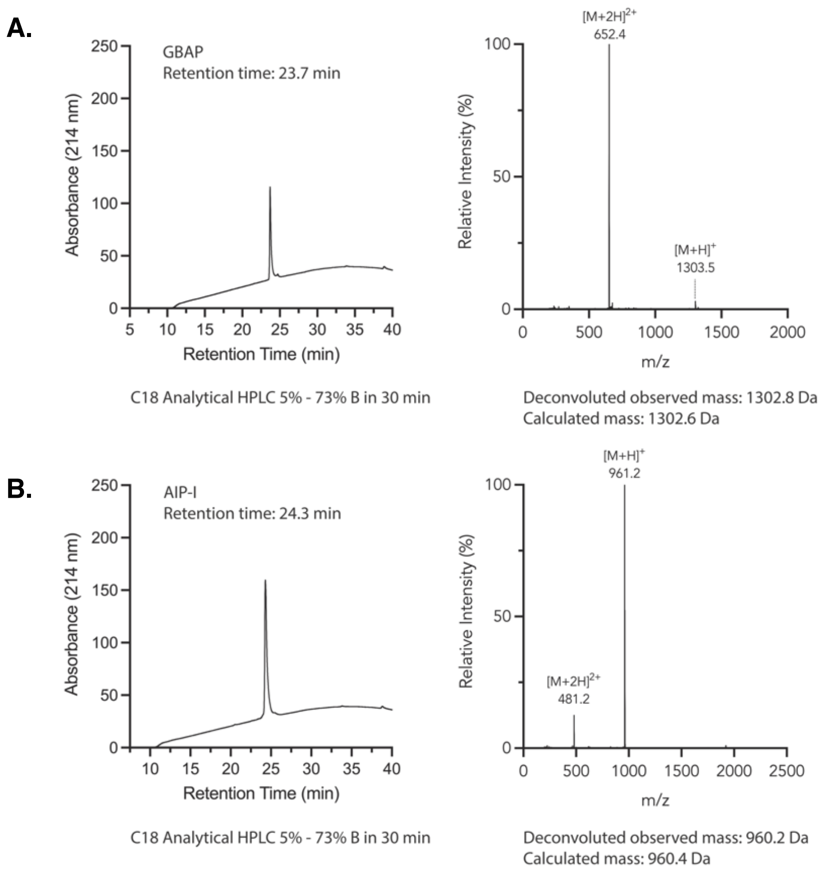


**Fig. S4. Characterization of synthetic autoinducing peptides. (A)** RP-HPLC and ESI-MS of GBAP. (B) RP-HPLC and ESI-MS of AIP-1

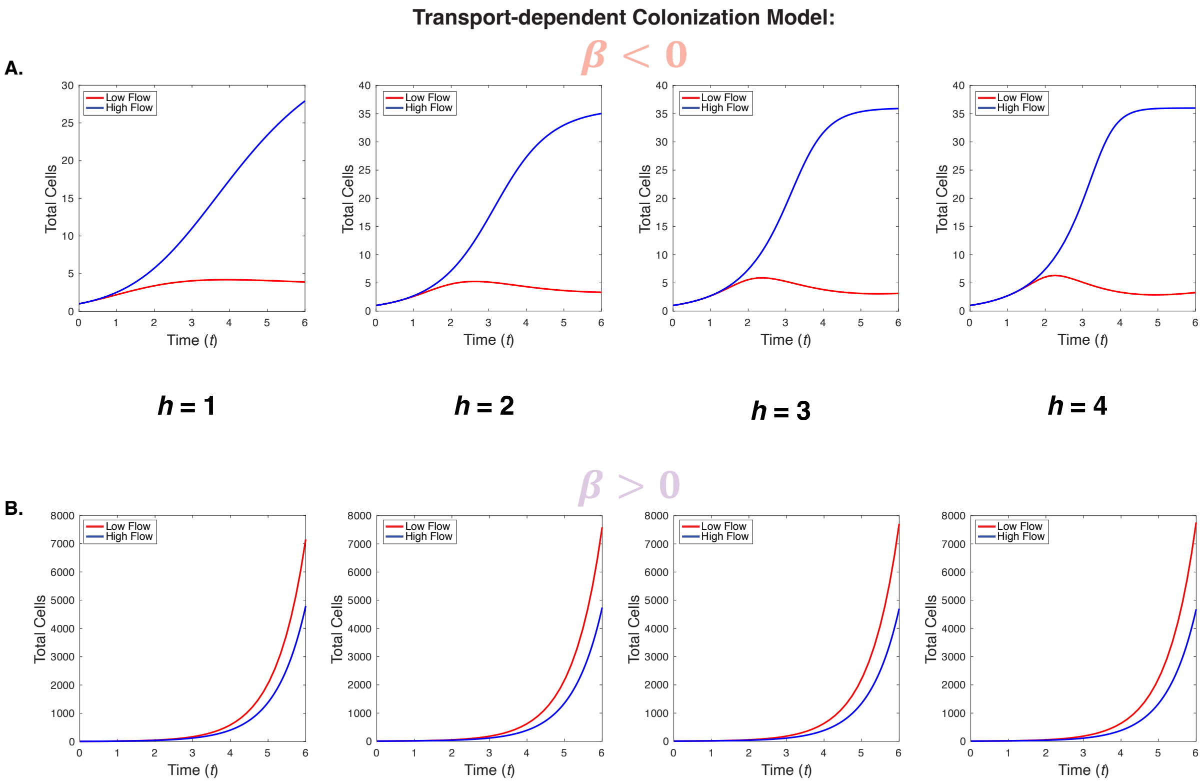


**Fig. S5. Varying the Hill coefficient for the transport-dependent colonization model (A)** Results for our ODE transport model using a range of Hill coefficients (h=1-4, left to right) for a positive β term. In all simulations, we observe larger final cell populations in the high flow condition, compared to the low flow condition. (B) Results for our ODE transport model using over a range of Hill coefficients (h=1-4, left to right) for a negative $\beta$ term. Once again, the qualitative behavior is consistent across all choices of h, indicating the insensitivity to the specific value chosen for the Hill coefficient. For all modeling in Fig. 2, we used h=3.

**Transport-dependent colonization model: Parameters**

| | $\beta < 0$ | $\beta > 0$ |
|---|---|---|
| $g$ | 0.5 | 0.8 |
| $\beta$ | -1 | 1 |
| $h$ | 3 | 3 |
| $K$ | 45 | 45 |
| $p$ | 5 | 5 |
| $q$ | [0.4, 4] | [4,40] |
| $c_o$ | 0 | 0 |
| $n_o$ | 1 | 5 |

**Table S1. Parameters used for the transport-dependent colonization model.** We note that the rates of attachment of new cells (a) and quorum sensing-independent detachment (r) as well as diffusion away from the colony (D) are set to 0.

**Mechanics-dependent colonization model: Parameters**

| | Parameter Values |
|---|---|
| $k_s$ | 1 |
| $l_0$ | 1 |
| $k_b$ | $1x10^{-3}$ |
| $d$ | 1 |
| $f$ | [0.001, 0.01] |

**Table S2. Parameters used for the mechanics-dependent colonization model.**

**Mechanics-dependent colonization model: Probabilities**

| | **Probability** | **Variables and Constants** |
|---|---|---|
| **Cell attachment** | $ae^{-\lambda y}$ | $a$ =0.2, $\lambda = 3$,<br>$y$ = height of cell above surface |
| **Cell detachment** | $ue^{-\alpha+m}$ | $u$ =1, $\alpha = 5$,<br>$m$ = height of neighbor cells |
| **Origin detachment** | $1 \times 10^{-6}$ | |
| **Chain breakage**<br>-Check for breakage if spring tension ($\tau$) > 0.02<br>- For *ΔatlA* mutant chains, check breakage if spring tension ($\tau$) > 0.022 | $1 - be^{-\omega\tau}$ | $b$ = 1, $\omega = 0.01$,<br>$\tau$ = spring tension |

**Table S3. Probabilities used for the mechanics-dependent colonization model.**

**Supplemental Movies**

**Movie S1.** *E. coli* low flow response. Representative *E. coli* experimental run in the low flow condition. Experiments ran for six hours and images were taken every five minutes.

**Movie S2.** *E. coli* high flow response. Representative *E. coli* experimental run in the high flow condition. Experiments ran for six hours and images were taken every five minutes.

**Movie S3.** *S. pneumoniae* low flow response. Representative *S. pneumoniae* experimental run in the low flow condition. Experiments ran for six hours and images were taken every five minutes.

**Movie S4.** *S. pneumoniae* high flow response. Representative *S. pneumoniae* experimental run in the high flow condition. Experiments ran for six hours and images were taken every five minutes.

**Movie S5.** *S. aureus* low flow response. Representative *S. aureus* experimental run in the low flow condition. Experiments ran for six hours and images were taken every five minutes.

**Movie S6.** *S. aureus* high flow response. Representative *S. aureus* experimental run in the high flow condition. Experiments ran for six hours and images were taken every five minutes.

**Movie S7.** *E. faecalis* low flow response. Representative *E. faecalis* experimental run in the low flow condition. Experiments ran for six hours and images were taken every five minutes.

**Movie S8.** *E. faecalis* high flow response. Representative *E. faecalis* experimental run in the high flow condition. Experiments ran for six hours and images were taken every five minutes.

**Movie S9.** Surface colonization of a co-culture of *P. aeruginosa* and *S. aureus* in minimal flow. Representative movie of *P. aeruginosa* (red fluorescence) and *S. aureus* (green fluorescence) co-culture in the minimal flow condition (shear rate 2/s) for the standard six hour experimental run.

**Movie S10.** Surface colonization of a co-culture of *P. aeruginosa* and *S. aureus* in high flow. Representative movie of *P. aeruginosa* (red fluorescence) and *S. aureus* (green fluorescence) co-culture in the high flow condition for the standard six hour experimental run.

**Movie S11.** *E. faecalis* breakage event in low flow (example 1). An excerpt from the experimental low flow *E. faecalis* run, demonstrating a breakage event of a chain.

**Movie S12.** *E. faecalis* breakage event in low flow (example 2). A second excerpt from the experimental low flow *E. faecalis* run, demonstrating a breakage event of a chain.

**Movie S13.** *E. faecalis* breakage event in high flow (example 1). An excerpt from the experimental high flow *E. faecalis* run, demonstrating a breakage event of a chain.

**Movie S14.** *E. faecalis* breakage event in high flow (example 2). A second excerpt from the experimental high flow *E. faecalis* run, demonstrating a breakage event of a chain.

**Movie S15.** *S. aureus* spontaneous detachment or attachment. Representative movie of a low flow experiment of *S. aureus* with a minimal media with no glucose to observe the spontaneous detachment and attachment (from upstream attachment).
